# Multi-omic characterization of human sural nerves across polyneuropathies

**DOI:** 10.1101/2024.12.05.627043

**Authors:** Michael Heming, Jolien Wolbert, Anna-Lena Börsch, Christian Thomas, Anne K. Mausberg, I-Na Lu, Julia Tietz, Finja Dienhart, Christine Dambietz, Anne-Kathrin Uerschels, Kathy Keyvani, Fabian Szepanowski, Bianca Eggert, Kai C. Liebig, Kathrin Doppler, Nurcan Üçeyler, Julieta Aprea, Andreas Dahl, Ruth Stassart, Robert Fledrich, Heinz Wiendl, Claudia Sommer, Mark Stettner, Gerd Meyer zu Hörste

**Author notes:** These authors contributed equally (Michael Heming, Jolien Wolbert).

## Abstract

Diseases of peripheral nerves termed polyneuropathies (PNPs) are common, mechanistically heterogeneous, and challenging to diagnose. Here, we integrated single nuclei transcriptomics of peripheral nerves from 33 human PNP patients and four controls (365,708 nuclei) with subcellular spatial transcriptomics. We identified novel and human-specific nerve cell type markers including unexpectedly heterogeneous perineurial fibroblasts. All PNPs shared a loss of myelinating and an increase in repair Schwann cells and endoneurial lipid-associated macrophages. Transcriptional changes affected multiple cells outside of the endoneurium across PNPs, suggesting PNPs as ‘pan-nerve diseases’. Spatially, PNPs showed a previously unknown perineurial hyperplasia and fibrotic dispersion and this was most pronounced in immune-mediated PNPs. Single cell transcriptomics supported the differential diagnosis of PNPs with potential for future unbiased diagnostic classification.

**One-sentence summary:** The first large-scale integrated single cell and spatial transcriptomic characterization of human peripheral nerves identifies novel cell markers and unexpected heterogeneity of perineurial cells, reveals polyneuropathies as ‘pan-nerve diseases’, and shows that single cell transcriptomics hold potential for unbiased nerve disease classification.

## Introduction

Polyneuropathies (PNPs) denote diseases of multiple peripheral nerves and range among the most common neurological diseases (*1*). Patients affected by PNPs suffer from progressive sensory and motor impairment with high socio-economic impact(*2, 3*). The underlying causes of PNPs are very diverse(*4–6*). Despite comprehensive diagnostic work-up, the underlying cause of PNPs often remains unclear (up to 20-30%) (*5*). Biopsy of the sensory sural nerve at the lateral ankle is often the final diagnostic step, but even biopsy does not lead to a diagnosis in many patients (*7*). Fully exploiting this precious biomaterial for mechanistic understanding and diagnostic potential is especially important to detect treatable causes such as immune mediated neuropathies, which account for up to 10% of the PNPs(*4, 5*). We and others previously performed single cell transcriptomic analyses of rodent peripheral nerves(*8–13*). However, human peripheral nerves have not been characterized using similar technologies.

Here, we provide the first large-scale single cell atlas of human peripheral nerves integrated with spatial transcriptomics. In addition to identifying novel and species-specific cell type markers, we detected a unique cellular composition of the perineurium and abundant nerve-associated leukocytes. In diseases, repair and damage Schwann cell (SC), and lipid-associated macrophages were shared across PNP etiologies. Multiple PNPs also affected cells beyond the endoneurium, causing perineurial cell proliferation and perineurial thickening preferentially in inflammatory PNPs. Finally, patients could be categorized into patient clusters identified solely by single cell transcriptomics. We thus discover potential for an unbiased diagnostic classification of PNPs.

## Results

### Comprehensive species-specific single nuclei transcriptional atlas of human sensory nerves

We first created a large single cell transcriptomics atlas of human sural nerves of 37 donors (Fig. 1A). Sural nerves were collected in three centers from 33 PNP patients and four controls (Suppl. Fig. 1A; Suppl. Tab. 1). Control samples (CTRL) were residual material of surgical sural nerve autografts (‘interpositions’) from patients with traumatic nerve injuries. We performed single nuclei RNA-sequencing (snRNA-seq) of all samples (Methods, Suppl. Fig. 1B, Suppl. Tab 2). After analytical removal of doublets and low quality nuclei and batch correction, this resulted in 365,708 total high-quality nuclei (Suppl. Fig. 1C-D, Suppl. Tab. 2). We then clustered the nuclei (henceforth termed ‘cells’) and annotated the clusters using predefined marker genes (Suppl. Fig. 2A, Suppl. Tab. 3) and automatic rodent-based annotation(*9, 10, 14*) (Suppl. Fig. 2B). We identified Schwann cells (*S100B, SOX10*), of both myelinating (mySC; *MPZ, MBP, PRX*) and non-myelinating (nmSC; *NCAM1*, *L1CAM, CDH2*) type and endoneurial fibroblast cells (endoC; *SOX9, PLXDC1, ABCA9*) (Fig. 1B, Supp. Fig. 2A). Schwann cells expressing features of damage (damageSC; *EGR1, FOS, JUN*) and of repair (*15*) (repairSC; *NGFR*, *ATF3, GDNF, RUNX2*) were novel compared to rodent data (*9, 10*). Moreover, we found epineurial (epiC; *CCBE1, COMP*) and perineurial fibroblast cells (periC1-3; *SLC2A1*/GLUT1, *KRT19, CLDN1*) (Fig. 1B, Supp. Fig. 2A). This perineurial cell heterogeneity was greater than previously described in rodents(*8–10, 12*). The periC3 cluster was distinguished from other pericyte clusters by genes associated with extracellular matrix and collagen fibril organization (Suppl. Fig. 2B). Vascular cells included vascular smooth muscle cells (VSMC: *ACTA2*, *CARMN*), pericytes (PC1-2; *PDGRFB, RGS5*) and endothelial cells (EC; *EGFL7, PECAM1*) (Supp. Fig. 2A). Based on known markers (Suppl. Fig. 2A) and published data(*14*) (Suppl. Fig. 2C) endothelial cells (EC) separated into lymphatic endothelial cells (LEC; *PROX1, LYVE1, FLT4*) and a venous (ven_EC: *PLVAP*, *ACKR1*) to capillary (capEC: *ABCG2*, *MFSD2A*) to arterial (artEC: *SEMA3G*, *HEY1*, *GJA5*) continuum. We identified a cluster of venous/capillary EC (ven_cap_EC2), which expressed blood-nerve barrier (BNB) markers *ABCB1* and *SLC1A1*(*16*), blood-brain barrier marker *MFSD2A*(*17*), and tight junction transcript *GJA1* and therefore likely represented EC of the BNB (Fig. 1B, Supp. Fig. 2A). Nerve-associated leukocytes were mainly of myeloid lineage outnumbering T/NK cells and B cells (Fig. 1B, Supp. Fig. 2A). We thus created a transcriptional cellular atlas of human peripheral nerves extending rodent studies(*8–13*).

**Figure 1:**
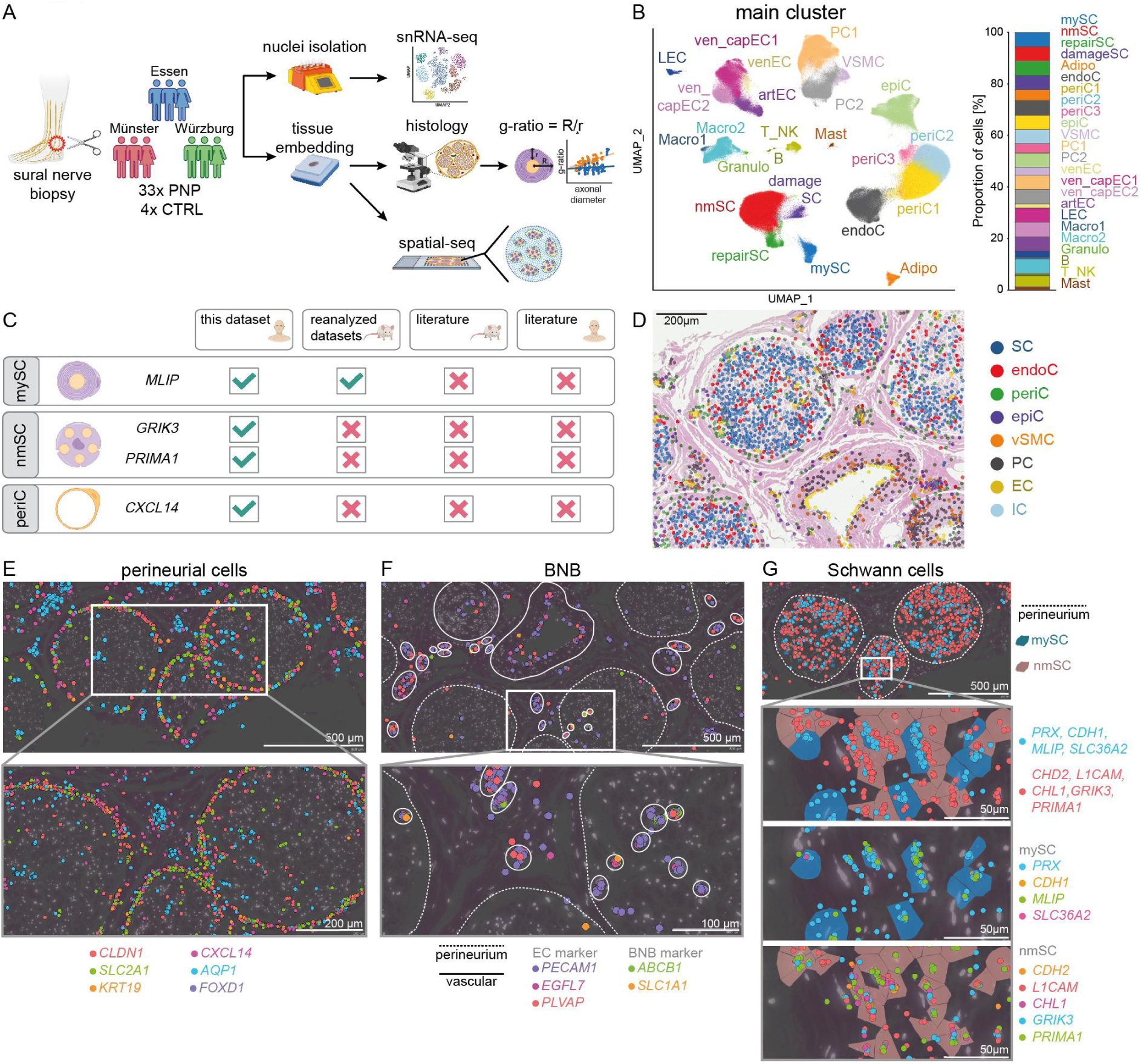
Defining the species-specific cellular landscape of human peripheral nerves. **(A)** Schematic overview of the experimental procedure. Sural nerve biopsies were collected from 33 polyneuropathy (PNP) patients and four controls (CTRL) from three German centers (Essen, Münster, Würzburg) and processed by single nuclei RNA-sequencing (snRNA-seq). Additionally, subcellular spatial transcriptomics (Xenium) was performed in a subgroup of eight patients. All 37 tissues were histologically characterized by quantifying myelin thickness (g-ratio), number of intactly myelinated axons and axon diameters. **(B)** UMAP of 365,708 high-quality nuclei from 37 human sural nerves, showing 23 main clusters. **(C)** Schematic depicting if novel marker genes of human myelinating Schwann cells (mySC), non-myelinating Schwann cells (nmSC), and perineurial cells (periC) found in this dataset were detected when reanalyzing published rodent dataset (Yim et al., Gerber et al., Wolbert et al., Supp. Tabl. 4-6) and whether these markers were previously described in rodent or human literature. **(D)** Representative section of an H&E staining overlaid with spatial-seq (Xenium) showing predicted clusters in spatial-seq of a sural nerve cross section of CTRL patient S24. Each dot represents one cell **(E)-(G)** Representative spatial-seq images of selected genes expressed in (E) perineurial cells, (F) in cells of the blood nerve barrier (BNB), and (G) in mySC (blue highlight) and nmSC (red highlight) in the sural nerve of CTRL patient S24. Each dot represents the expression of one transcript, a dotted line marks the perineurium, and a solid line surrounds individual vessels.

We systematically compared cell type markers in humans with their published rodent counterparts(*8–10*) (Suppl. Tab. 4-6). As expected, the majority of transcripts were shared between species. However, several genes expressed in mySC (*MLIP*), nmSC (*GRIK3*, *PRIMA1*), and periC (*CXCL14*) were undescribed in rodent or human literature. When re-analyzing rodent datasets(*8–10*), we detected *Mlip* in mySC in two studies(*9, 10*); albeit previously unmentioned (Fig. 1C, Suppl. Tab. 4-5). In contrast, *GRIK3*, *PRIMA1* and *CXCL14*, were not detectable in rodents (Suppl. Fig. 2D). We thus identified novel and human-specific transcripts across nerve-associated cells.

### Spatially confirming novel transcriptional cell type markers

For morphological confirmation, we employed *in situ* amplification-based spatial transcriptomics with subcellular resolution (‘spatial-seq’)(*18*). A custom panel of 99 combined known (e.g., *PRX*) and novel (e.g., *GRIK3*) transcripts (Suppl. Tab. 7) while excluding highly transcribed genes to prevent optical overcrowding (Methods). Visualizing these targets in a formalin-fixed paraffin-embedded (FFPE) cross-section of a human sural nerve graft (CTRL) replicated the overall cellular organization. We matched each cell in spatial-seq to a snRNA-seq cluster, aggregated these into 8 larger groups and then visualized them (Fig. 1D). As expected, the nerve consisted of multiple fascicles formed by endoneurial cells (SC, endoC) surrounded by perineurial cells (periC). Vascular cell types formed vessels of their respective morphology (Fig. 1D). Epineurial cells (epiC) were located in between fascicles and were interspersed with immune cells that were rare in the endoneurial areas (Fig. 1D). We thus provide an unbiased histological annotation of peripheral nerve cells and clusters.

Previously undescribed perineurial markers (*CXCL14*) indeed co-localized with known perineurial fibroblast transcripts (*CLDN1, SLC2A1, KRT19*)(*11*) in the perineurium (Fig. 1E). The BNB markers *ABCB1* and *SLC1A1,* expressed by the ven_capEC2 cluster in snRNA-seq, were enriched in endoneurial vessels together with known EC markers (*PECAM1, EGFL7*)(*11, 19*) (Fig. 1F). This supports that the ven_capEC2 cluster represents BNB cells. Other vascular cell markers (e.g., *SEMA3G* in artEC) were associated with their respective vessel type (Suppl. Fig. 3A). Known pan-SC markers (*SOX10, S100B, EGR2*)(*11*) were expressed across the endoneurium (Suppl. Fig. 3B). Known markers of mySC (*PRX, CDH1, SLC36A2*)(*9, 13*) co-localized with *MLIP* identified in the mySC cluster (Fig. 1G.). Markers of nmSC (*L1CAM, CDH2, CHL1*)(*12, 19, 20*) co-localized with novel transcripts of the nmSC cluster (*PRIMA1, GRIK3*) (Fig. 1G). We thus spatially validated novel human markers of perineurial fibroblasts, the BNB, and Schwann cells.

### Human peripheral nerves contain complex immune cells

We next deeply sub-clustered all immune cell (IC) nuclei (n = 18,436) at high resolution (Fig. 2A). Megakaryocytes/platelets (*PF4, GP9, PPBP*) and red blood cells (*HBA, HBB*) were undetected arguing against blood contamination (Fig. 2A). The majority of IC clusters (79% of all IC nuclei) were of myeloid lineage; mainly macrophages (Macro1-18: *LYZ*, *CD14*, *MRC1*, *CD163*, *MS4A7*) (Fig. 2A, Suppl. Fig. 4A, Suppl. Tab. 8). The phenotype and ontogeny of nerve-associated macrophages differ depending on their endo- vs. epineurial location in rodents (*8, 21, 22*). In humans, macrophage clusters (*MS4A7*) also exhibited a gradient of expression from endoneurial (*CX3CR1, TREM2)* to epineurial (*LYVE1, FOLR2, TIMD4*) markers (Fig. 2B). One representative tentatively endoneurial macrophage cluster (Macro18) was enriched in lipid metabolism-related pathways (Fig. 2C). Markers of epi- (*FOLR2*) vs. endoneurial (*CX3CR1*) macrophages (*MS4A7*)(*21*) were preferentially detected in their epi- vs. endoneurial location (Fig. 2D). We additionally identified smaller myeloid clusters that could be assigned to classical dendritic cells (DC) type 1 (cDC1: *CLEC9A*, *XCR1*, *BATF3*), plasmacytoid DC (pDC: *CLEC4C*, *IRF8*), and classical DC type 2 (cDC2: *FCER1A*, *CD1C*, *CLEC10A*) (Fig. 2A, Suppl. Fig. 4A). cDC2 have not been previously described in nerves(*13*). The T/NK cell clusters ranged from naive CD4 (*CCR7*, *SELL/*CD62L), Treg (*FOXP3*), MAIT (*KLRB1*, *CXCR6*) to CD8 T Cells (*CD8A)* and NK cells (*FCGR3A*, *NKG7*) (Fig. 2A, Suppl. Fig. 4A). Naive B cells (*CD19*, *MS4A1/*CD20, *CD79B,*) and plasma cells (*JCHAIN*, *SDC1*/CD138) (Fig. 2A, Suppl. Fig. 4A) were detected and mainly expressed *IGHG1*, *IGHG3*, *IGHG4*, and *IGHA1* heavy chain genes (Fig. 2E). We thus delineated nerve-associated leukocytes in humans in unprecedented detail.

**Figure 2:**
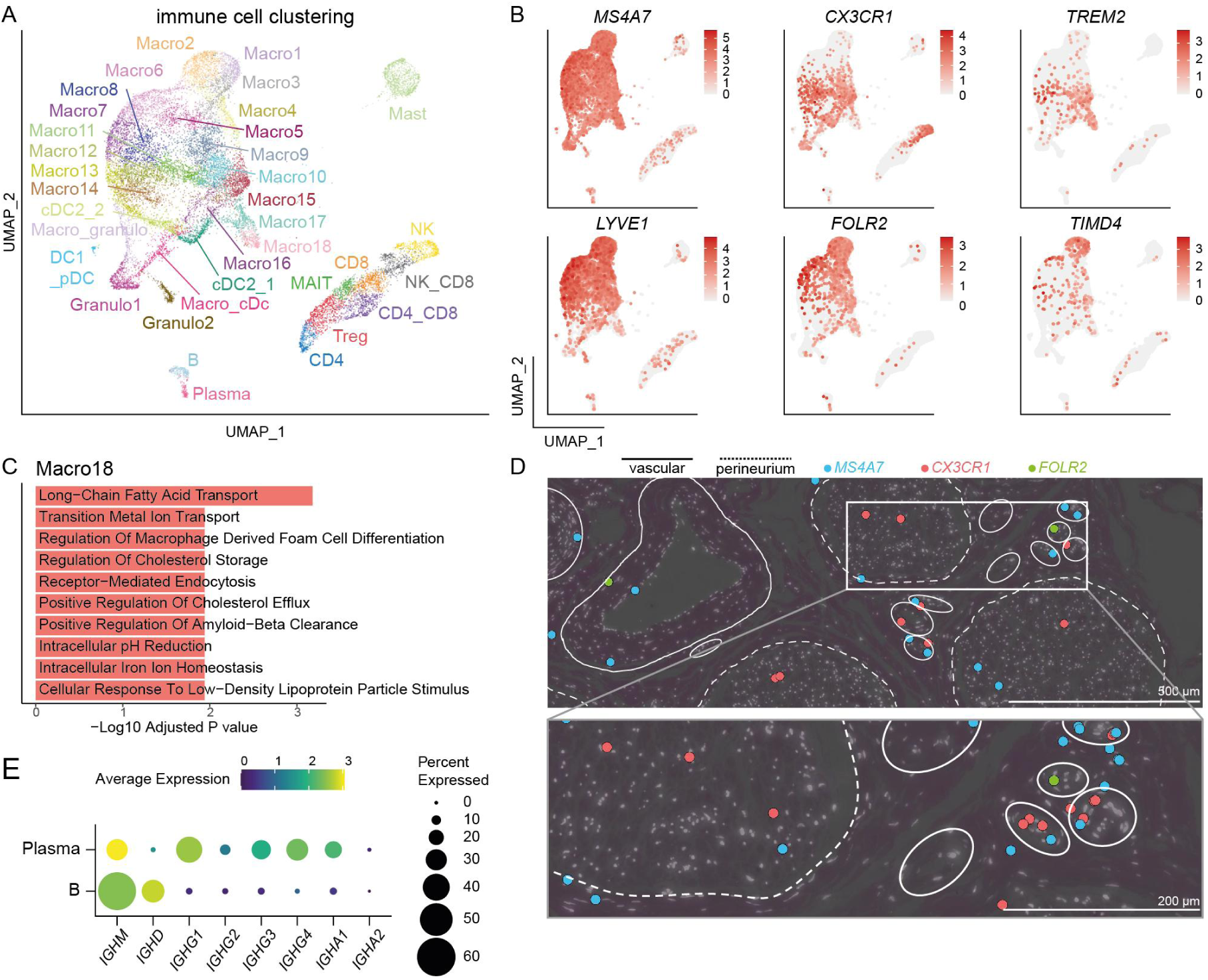
Heterogeneity of human nerve-associated immune cells. **(A)** UMAP of 18,436 nuclei representing 35 immune cell (IC) subclusters of 37 human sural nerves. **(B)** Feature plots of known endoneurial and epineurial macrophage marker genes. Color encodes gene expression. **(C)** Gene ontology term enrichment analysis of marker genes (log_2_ fold change > 2, adjusted p value < 0.001) expressed by the Macro18 IC cluster in a one vs. all IC cluster comparison. **(D)** Representative spatial-seq images of the macrophage marker *MS4A7* and the endoneurial macrophage markers *CX3CR1* and *TREM2* in the sural nerve of CTRL patient S24. Each dot represents the expression of one transcript, a dotted line marks the perineurium, and a solid line surrounds individual vessels. **(E)** Expression of immunoglobulin heavy (*IGH*) chain genes in the B cell (B) and plasma cell (plasma) cluster of the IC subclusters. Dot size encodes the percentage of cells expressing the transcript, and color shows the average gene expression.

### Non-endoneurial cell types profoundly respond to polyneuropathy

We characterized how PNP affected peripheral nerves (Suppl. Fig. 1E). The most apparent change in PNP was a loss of mySC and the occurrence of damageSC and repairSC (Fig. 3A), induced by nerve damage in rodents(*15, 23–25*). Among IC subclusters, an increase in the endoneurial lipid-associated Macro18 subcluster was the most abundant change in PNP patients (Fig. 3A). Notably, the periC3 cluster consistently increased in PNP (Fig. 3A). When testing which *individual* single nuclei were associated with disease, the PNP status positively correlated with nuclei in the leukocyte and repairSC clusters and negatively correlated with the mySC cluster after correcting for confounders (Methods) (Fig. 3B, left panel). This means that mySC are less likely to be present in PNP patients on a per nucleus level. In addition, the periC3 cluster positively correlated with PNP disease status (Fig. 3B, left panel). These clusters similarly correlated with clinical disease severity as quantified by the clinical INCAT score (Fig. 3B, middle panel) and myelin thickness (inversely measured by g-ratio) (Fig. 3B, right panel). PNPs thus induced profound compositional changes that correlated with disease severity and known measures of PNP and specifically affected perineurial cells.

**Figure 3:**
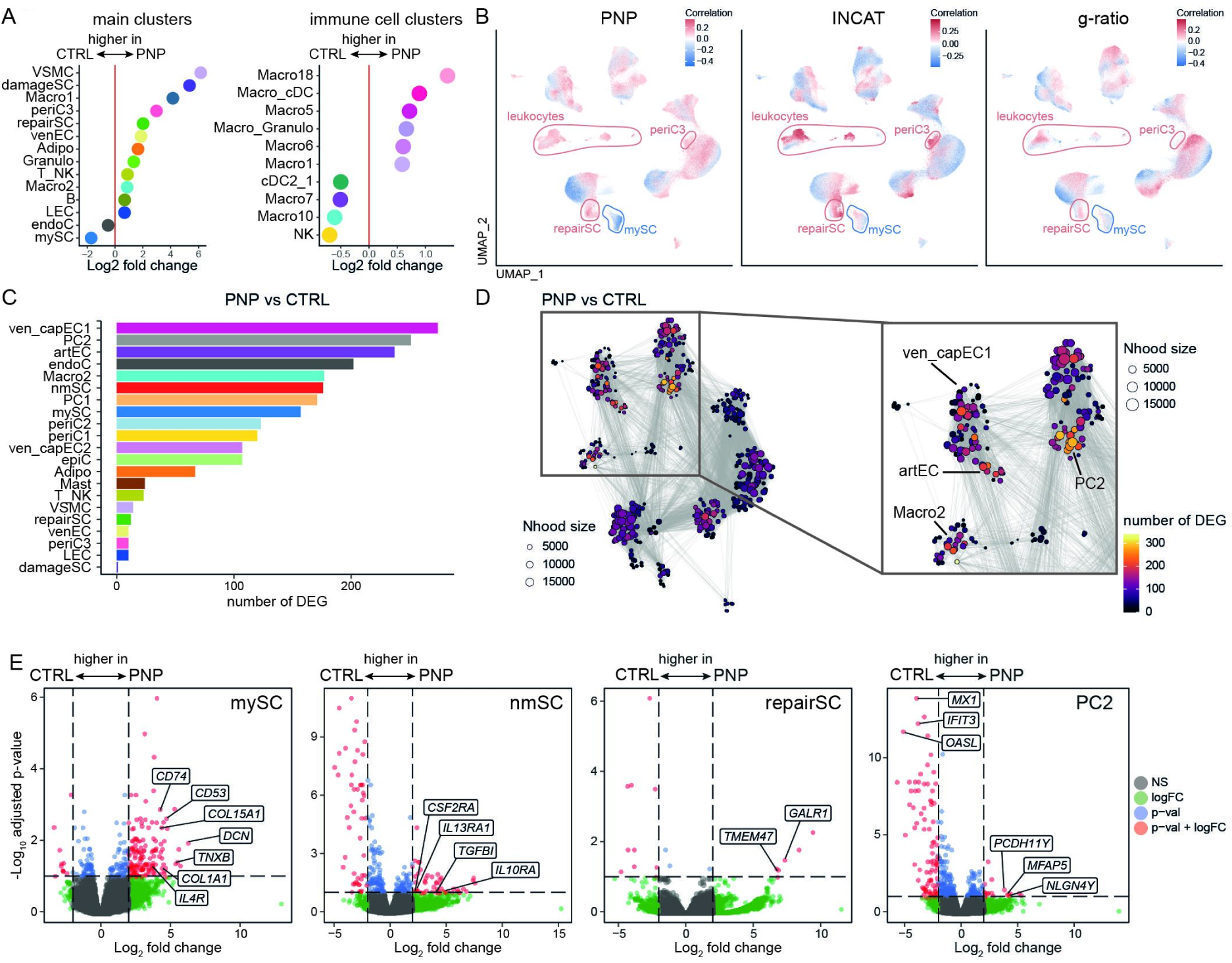
Identifying a pan-neuropathy cellular transcriptional response pattern. **(A)** Differences in cluster abundance in patients with polyneuropathy (PNP) vs. controls (CTRL), left: main clusters (Fig. 1B), right: immune cell subclusters (Fig.2A). Only clusters with a log_2_ fold change > 0.5 are shown. **(B)** Correlations between individual single nuclei and PNP status (right), clinical Inflammatory Neuropathy Cause and Treatment (INCAT) disability score (middle), and g-ratio of myelinated axons (right) are shown. Red indicates a positive correlation and blue a negative correlation. **(C)** The number of differentially expressed genes (DEG) per cell cluster were determined in a pseudo bulk approach and are visualized. **(D)** Neighborhood graph displaying the number of DEG between PNP and CTRL in a clustering-independent cellular neighborhood-based approach. Each dot represents a neighborhood, and the dot size represents the size of the neighborhood (Nhood), while edges depict the number of cells shared between neighborhoods. Color code shows the number of DEG. The right plot shows a magnification of the left plot, highlighting neighborhoods/clusters with the highest number of DEG. **(E)** Volcano plots of DEG in myelinating Schwann cells (mySC), non-myelinating SC (nmSC), repair SC, and pericytes 2 (PC2) between PNP and CTRL. The horizontal dashed line represents an adjusted p value of 0.1 and the vertical dashed lines display a log_2_ fold change of 2 Selected transcripts above those thresholds are labeled.

We then characterized which and how nerve cells *transcriptionally* responded to disease. The number of differentially expressed genes (DEG) was highest in vascular clusters (ven_capEC1, PC2, artEC) followed by endoneurial fibroblasts (endoC) (Fig. 3C). Using a clustering-independent cellular neighborhood-based approach, we accordingly found that neighborhoods with most DEG were located in vascular clusters (PC2, ven_capEC1, artEC) and in the Macro2 cluster (Fig. 3D). Next, we more specifically tested how PNP influenced gene expression of selected nerve-associated cells (mySC, nmSC, repairSC, PC2) (Fig. 3E). We found that many of the DEG in the mySC cluster were associated with fibrotic tissue remodeling (e.g., *DCN, TNXB*), extracellular matrix formation (e.g., *COL1A1, COL15A1*), and immune regulation (e.g., *CD53, IL4R, CD74*). The repairSC cluster expressed genes for nerve repair and neuronal differentiation (*GALR1, TMEM47*) (Fig. 3E; Suppl. Tab. 9). GO term enrichment analysis showed that upregulated genes were associated with cell differentiation in the mySC cluster and with cell migration in the nmSC cluster (Suppl. Fig 4B). In summary, PNPs induced transcriptional changes in Schwann cells but also widely outside of the endoneurium, especially in non-endoneurial vascular cells. PNPs thus surprisingly affect the cellular micro-milieu of peripheral nerves beyond the endoneurium and may thus constitute ‘pan-nerve diseases’.

### PNP subtypes preferentially affect different cellular compartments of peripheral nerves

The heterogeneity of PNP mechanisms is poorly defined. We classified the available PNP patients by integrating all diagnostic information (Suppl. Tab. 1) into seven subtypes of PNPs: vasculitic (VN, n = 5), chronic inflammatory demyelinating (CIDP, n = 9), chronic idiopathic axonal (CIAP, n = 11), cancer-associated paraproteinemic (PPN, n = 2), diabetic (DPN, n = 2), other inflammatory (OIN, n = 2), and other non-inflammatory (ONIN, n = 2) (Fig. 4A). Histological characterization (Supppl. Fig. 5) identified that myelin thickness (Suppl. Fig. 5B-C) and the average number of intactly myelinated axons (Suppl. Fig. 5D) decreased in PNPs, while axon diameter was less affected (Suppl. Fig. 5E). Histological findings in PNP subtypes were in accordance with expectations and correlated with electrophysiology (Suppl. Fig. 5F) and with cluster proportions determined by snRNA-seq (Suppl. Fig. 5G).

**Figure 4:**
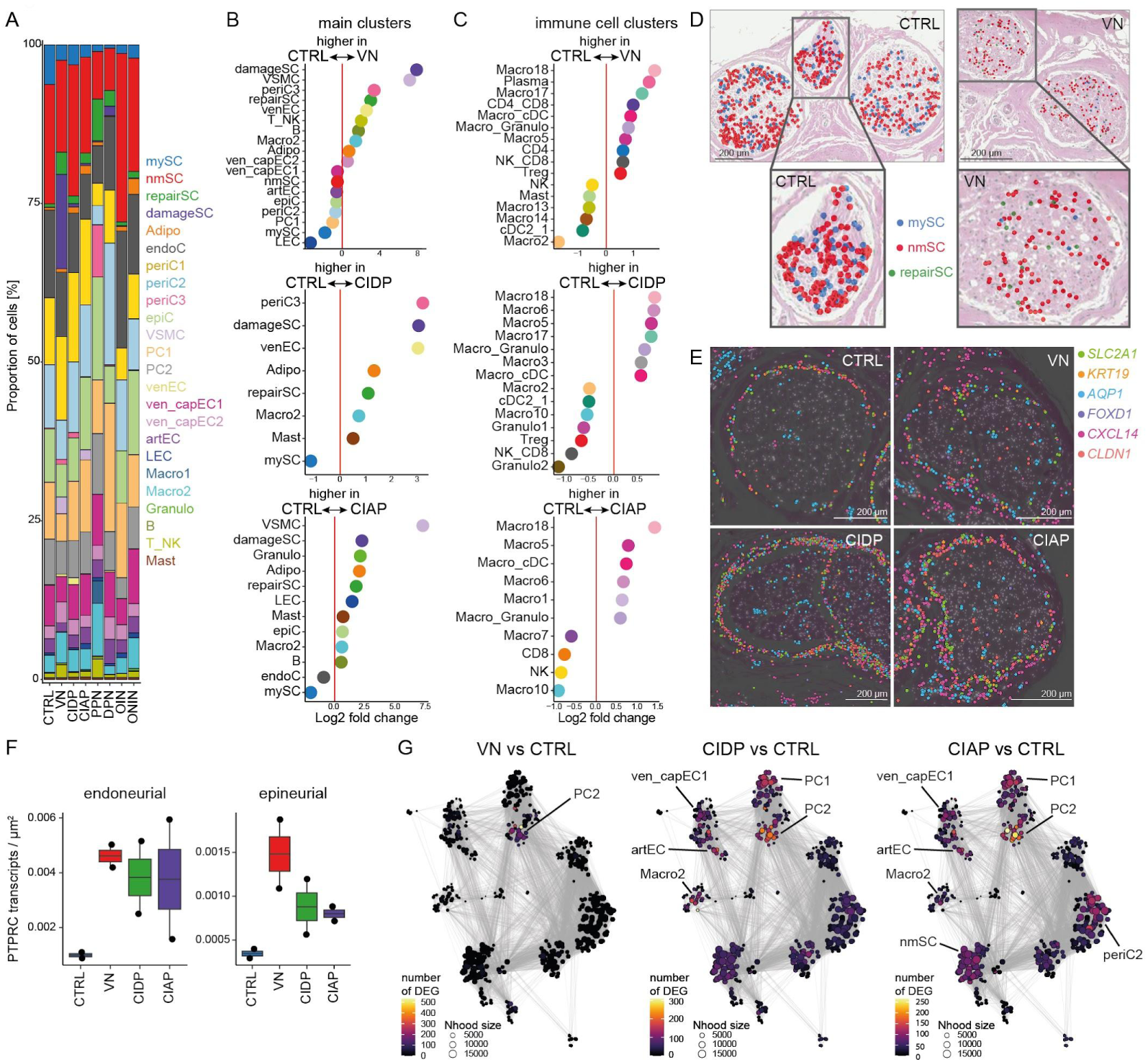
PNP subtypes affect different cellular compartments of peripheral nerves. **(A)** Proportion of cells split by disease group and colored by the main clusters. **(B) - (C)** Changes of cluster abundance in patients with vasculitic (VN), chronic inflammatory demyelinating (CIDP), and chronic idiopathic axonal (CIAP) polyneuropathy vs. controls (CTRL). The left plots show changes of the main clusters (as in Fig. 1B), the left plots show differences in the immune cell subclusters (Fig. 2A).Only clusters with a log_2_ fold change > 0.5 are shown. **(D)** Representative sections of H&E staining of sural nerves overlaid with spatial seq (Xenium) showing predicted Schwann cell clusters in CTRL (S24, left plot), and VN (S30, right plot) patients. Each dot represents one cell. **(E)** Representative spatial transcriptomics images of perineurial marker genes of CTRL (S24), VN (S30), CIDP (S01), and CIAP (S14). Each dot represents the expression of one transcript. **(F)** The density of *PTPRC* transcripts in the endoneurial (left pot) and epineurial (right plot) per disease group (n = 2 per disease group). **(G)** Neighborhood graphs displaying the number of DEG between PNP patients with VN (left), CIDP (middle) or CIAP (right) and CTRL in a clustering-independent cellular neighborhood-based approach. Dot size represents neighborhoods (Nhood), while edges depict the number of cells shared between neighborhoods. Color code shows the number of DEG.

**Figure 5:**
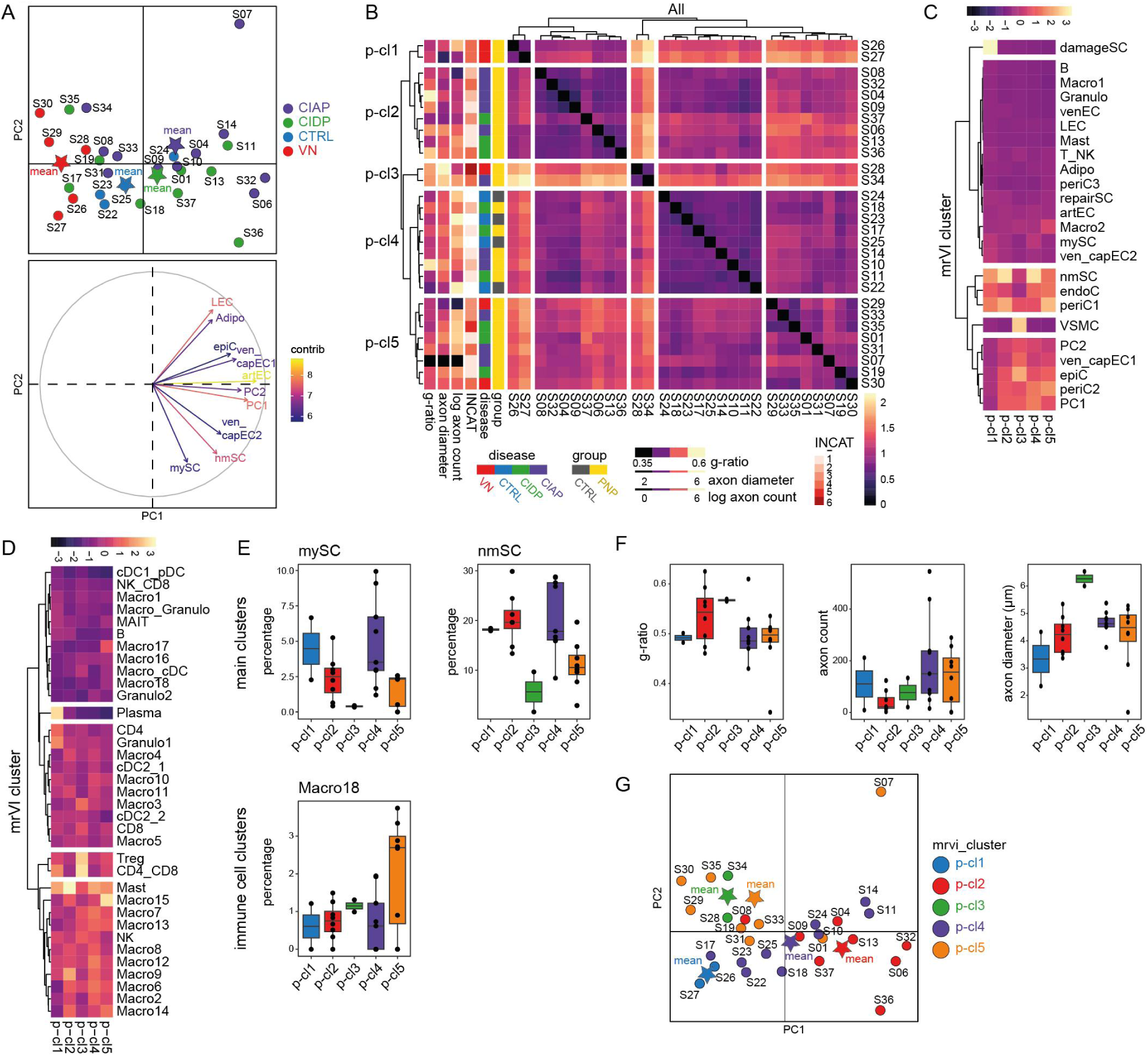
Hypothesis-free PNP patient classification. **(A)** Upper plot: PCA based on the relative cluster abundance in 29 patients categorized by disease group (vasculitic PNP, VN, n = 5), chronic inflammatory demyelination (CIDP, n = 9), chronic idiopathic axonal (CIAP, n = 11)) and controls (CTRL, n = 4). Each dot represents a patient, the star represents the group mean. Lower plot: Individual variables of the PCA. The contribution of each variable is color-coded. **(B)** Patient-patient comparisons were conducted with a deep generative model (multi-resolution variational inference). Patient-patient distances were visualized in a hierarchically clustered heatmap. The heatmapis annotated with clinical (group, disease, INCAT disability score) and histological (number of intactly myelinated axons, g-ratio, axon diameter). **(C-D)** Scaled cellular abundance per patient clusters (defined in **B**) of main clusters (**C**) and immune cell subclusters (**D**) visualized in a clustered heatmap. **(E)** Abundance of selected cell clusters per patient clusters. **(F)** Histological measures (g-ratio, number of intactly myelinated axons, axon diameter) per patient cluster. (**G**) PCA based on the relative cluster abundance (as in A), but categorized by patient-clusters.

We next compared cluster proportions between groups with at least four samples per group (VN, CIDP, CIAP, CTRL), while disregarding others. In comparison with controls, a relative increase in damageSC and repairSC, and loss of mySC was detected in VN, CIDP, CIAP (Fig. 4B). A gain of the perineurial periC3 cluster and the venEC cluster was shared between VN and CIDP. An increase in the VSMC cluster was found in both VN and CIAP. Loss of the LEC cluster was specific to VN (Fig. 4B). The lipid-associated Macro18 cluster was increased in all three PNP groups compared to CTRL (Fig. 4C). Only VN showed an expansion of the plasma cluster. Each PNP subtype thus induced a unique pattern of compositional cellular changes.

We then analyzed spatial transcriptomics of sural nerves from VN, CIDP, and CIAP patients vs. controls (n = 2 per group). Predicted repairSC were increased in the endoneurium in VN (Fig. 4D). Predicted mySC were considerably less abundant in VN than in CTRL (Fig. 4D) and this was also true but less prominent for CIDP and CIAP (Suppl. Fig. 5H-I). Quantification of leukocyte-associated transcripts (*PTPRC*/CD45, *MS4A1*/CD20, *CD3E*) showed that the density of endo- and epineurial B and T cells increased in VN, CIDP and in CIAP compared to CTRL (Fig. 4F, Suppl. Fig. 5J-K). Predicted T_NK cells were more abundant in VN and less so in CIDP and CIAP than in CTRL (Suppl. Fig. 5 L,M). We thus spatially validated the loss of mySC and gain of repairSC and leukocytes in PNPs with most most prominent effects in VN.

We next aimed to spatially validate the increase in the periC3 (perineurial cell) cluster in snRNA-seq. Plotting known (e.g. *CLDN1, SLC2A1, KRT19*) and novel (e.g. *CXCL14,*) perineurial markers, identified the perineurium as a single cell layer in CTRL, but as 3-5 cell layers in PNP samples which was most pronounced in CIDP and appeared regionalized (Fig. 4E). The perineurium was also more dispersed and heterogeneous in PNP samples (Fig. 4E). Perineurial hyperplasia driven by expansion of PNP-specific perineurial cells thus occurs across PNPs, but preferentially in CIDP.

Next, we tested for subtype-specific transcriptional alterations. In VN, most DEG were located in neighborhoods in the PC2 (pericyte) cluster (Fig. 4G) in accordance with vascular-focused inflammation. In CIDP, DEG were also identified in neighborhoods of vascular clusters (ven_capEC1, artEC, PC2) and the Macro2 cluster. In CIAP, DEG were distributed in a more widespread pattern (Fig. 4G). Each PNP subtype thus induced unique compositional and transcriptional responses.

### Hypothesis-free PNP patient classification

Finally, we speculated that PNP patients could be classified purely based on biopsy snRNA-seq. Principal component analysis (PCA) based on relative cluster abundance especially separated VN from other patients and this was driven by vascular clusters (e.g. artEC, PC1) (Fig. 5A). CIDP and CIAP patients exhibited a larger dispersion potentially indicating a more heterogeneous pathogenesis (Fig. 5A). Integrating histological measures into the PCA analysis did not show evident enrichment of a certain histological feature in the PCA space.(Suppl. Fig. 6A) indicating subsetting beyond histology alone.

We next used the multi-resolution variational inference (MrVI) (*26*) approach to group patients. This deep generative model is designed for sample-sample comparisons using single cell information (Methods). Samples formed five patient clusters (p-cl; Fig. 5B). A ‘normal’ patient cluster of patients (p-cl4) included all CTRL samples and featured normal cluster composition, low disability and normal histological measures (Fig. 5B-F, Suppl. Fig. 6B). P-cl2 described as ‘large-axon-loss’ exhibited a reduction of mySC (Fig. 5E), moderate disability, strong reduction in intact axons, and thinly myelinated remaining axons (Fig. 5B,F, Suppl. Fig. 6B). P-cl5 showed severe disability, despite many intact axons and intact myelin thickness but reduction of the endoneurial mySC and nmSC clusters (Fig. 5B,E), and an increase in endoneurial lipid-associated Macro18 (Fig. 5E; Suppl. Fig. 6D), indicating immune-associated disability. P-cl1 was enriched in VN patients, showed severe disability, small axons, no relevant demyelination, and an increase in plasma cells (Fig. 5B, D,F) potentially indicating antibody-driven pathology. When plotting the mrVI-based patient classification into the cell cluster abundance-driven PCA, the mrVI patient clusters were separated to some extent (Fig. 5G). Single nuclei transcriptomics of sural nerves thus categorized PNP patients in a non-hypothesis-driven manner.

## Discussion

We aimed to exploit the full potential of human nerve biopsies using state-of-the-art techniques and present a single nuclei transcriptomics atlas comprising 365,708 nuclei of 33 PNP patients and four controls integrated with subcellular spatial transcriptomics. Capitalizing on this first application to human peripheral nerves and a dataset 10-fold larger than previous rodent studies(*8–13*) we discovered and validated previously undescribed transcriptional markers of perineurial fibroblasts (*CXCL14*) and myelinating (*MLIP*) and non-myelinating SC (*GRIK3, PRIMA1*), which were partially human-specific. In human PNP, we identified perineurial hyperplasia driven by a PNP-specific perineurial cell type. Transcriptional changes widely affected multiple non-endoneurial cell populations unexpectedly suggesting PNPs as ‘pan-nerve’ diseases. Using single cell transcriptomics, we identified a PNP subtype-specific composition and transcriptome with diagnostic potential. Additionally, single cell data showed potential for unbiased and potentially mechanistically-driven classification of PNPs in the future.

Among novel cell type transcripts, the *MLIP* gene encodes a muscular laminin-interacting protein, required to maintain muscular integrity (*27*) which could serve similar functions in SC. The *PRIMA1*-encoded protein organizes esterases at the neuromuscular junction in terminal SC (*28*), but has not been described in nmSC. *GRIK3* encodes a glutamate ionotropic receptor (*29*) and variants in GRIK3 have been associated with hereditary neuropathy (*30*).These genes may thus be involved in structural organization of SC in human peripheral nerves. Human CXCL14 (synonymously BRAK) is constitutively expressed by epithelial tissues and regulates microglial development in the brain(*31*) and neurovascular patterning in the eye(*32*). It is thus conceivable that the perineurium, regarded as an epithelial barrier (*33*), expresses *CXCL14* contributing to nerve organization. The species-specificity of some of the markers (*GRIK3, PRIMA1, CXCL14*) could be due to i) size or phylogeny of the mammal, ii) diseased samples dominating our dataset, or iii) the purely sensory sural nerve analyzed in humans. Discrepancies are unlikely to be driven by single nuclei vs. cell approaches because one rodent study analyzed nuclei(*9*) without detecting these markers.

The perineurium is a lamellated structure made up of concentric cell layers bordered on each side by a basement membrane(*34*). A linear relationship has been described between fascicle diameter and perineurium thickness(*35*), i.e. larger fascicles have a thicker perineurium. Additionally, distal sections have a greater perineurium thickness than proximal segments(*35*). Thickening of the perineurial basement membrane is a characteristic of diabetic PNP(*36*). Focal or generalized perineurial thickening occurs in ‘perineuritis’ - a rare inflammatory disease(*37*).Overall, perineurial thickening *per se* has not been studied systematically in PNPs. Proliferation of PNP-specific perineurial cells could reflect a maladaptive response of the nerve microstructure to chronic damage.

Diagnosing polyneuropathies can be challenging due to the diverse range of possible causes(*5*). Even histology of the sural nerves often fails to establish the etiology of PNP(*7*). Here, we used data-driven approaches to classify patients solely based on snRNA-seq into pathophysiological groups with distinct clinical phenotypes. We speculate that snRNA-seq could represent a PNP classification tool towards personalized neuro-pathology(*38–40*). Our study has limitations: Our patient cohort is large, but potentially confounded by immunosuppressive therapy in some patients (Suppl. Tab. 1) and it is unbalanced regarding disease duration, sex and center of origin. Even larger multi-centric multi-national funding, collection and analysis efforts will be required in the future. Sural nerve biopsies are invasive with some inherent risk and are usually performed in difficult to diagnose patients(*7*). Our patient cohort is thus biased towards rare PNP etiologies and, for example, includes only two diabetic PNP; the most common cause of PNPs(*1, 4, 6*). We mitigate this by focussing on groups with at least 4 patients. We believe that novel techniques to enrich rare cell types(*41*) could identify PNP-associated cellular patterns in more readily available tissues such as skin biopsies in the future. Whether less expensive and less labor-intensive methods, such as bulk transcriptomic or methylome analysis of peripheral nerves, could similarly be used remains to be determined. In summary, our data revisit the cellular and anatomical architecture of peripheral nerves in a human-specific fashion and define a perspective for developing intra- and inter-PNP classifiers towards personalized PNP diagnosis.

## Materials and Methods

### Collection of patient samples

We collected sural nerves from 33 PNP patients from three centers: University Hospital of Münster (10 patients), Essen (11 patients), and Würzburg (12 patients). Additionally, we obtained sural nerves from four control patients in Essen with traumatic nerve injuries, who received a sural nerve autograft as interposition, but were unaffected by polyneuropathy (Suppl. Tab. 1). Residual material from the sural nerve grafts were used for control samples in this study. In PNP patients, sural nerve biopsy was performed as part of the clinical diagnostic workup and a small portion of the nerve (approximately 0.2 cm) was used for this study. In Münster and Essen, the samples were immediately fresh-frozen after surgery in dry ice-cooled methylbutane and stored in liquid nitrogen (0 - 14 months, Suppl. Tab. 1) until nuclei extraction. In Würzburg, the samples were embedded into OCT embedding matrix (Roth) until nuclei extraction (33 - 77 months, Suppl. Tab. 1). In Essen and Münster samples were collected from newly recruited patients. In Würzburg, samples were collected from an existing biobank of OCT embedded cryo-preserved sural nerve biopsies collected between 2016 and 2020. All experiments were carried out in accordance with the Declaration of Helsinki and were approved by the local ethical committees in Münster (2018-719-f-S), Essen (21-10376-BO), and Würzburg (238/17; 15/19). All patients gave written informed consent to sample collection.

### Patient characteristics

Patients were characterized clinically regarding diagnosis, disease activity, therapy response, disease duration, immunosuppressive therapy, relevant secondary diagnosis, electrophysiological studies and CSF analysis outlined in Suppl. Tab. 1. Patients were diagnosed according to the EAN/PNS criteria 2021 for CIDP(*42*) and according to the PNS criteria for VN.

### Nuclei extraction and purification

For each patient, approximately 10 mg pieces (median of 13.9 mg) were cut on dry ice as starting material from the original samples. Nerve pieces were cut into smaller pieces and processed using the Miltenyi Biotec nuclei extraction protocol. In summary, samples were transferred to a gentleMACS C-tube (Cat.no. 130-093-237) containing 2 ml lysis buffer (nuclei extraction buffer (Cat. no. 130-128-024) + 0.2 U/µl RNase inhibitor (EO0381) and processed on a gentleMACS dissociator with program 4C_nuclei_1. After dissociation, the nuclei suspension was filtered through a pluriStrainer Mini 70-µm cell strainer (pluriSelect®) and washed in 200-400 µl resuspension buffer (PBS with 0.1% BSA and 0.2 U/μL RNase inhibitor (EO0381) depending on sample size and expected nuclei count. Nuclei suspension was then filtered through a pluriStrainer Mini 40-μm cell strainer (pluriSelect®), placed on a 1.5 ml DNA-LoBind tube (Merck). The nuclei suspension was stained with Trypan Blue to manually assess nuclei viability and count using a Fuchs-Rosenthal chamber. Equal volumes between samples were used for downstream application single nuclei RNA seq.

### Single nuclei RNA-sequencing and generation of count matrices

Single nuclei suspensions were loaded into a Chromium Next GEM Chip G and placed into the Chromium X Single Cell Controller and processed with Chromium Next GEM Single Cell 3’ Kit v3.1 reagents (all 10X Genomics). Sequencing was performed on Illumina Nextseq 2000 and Novaseq 6000 with a 28-8-0-91 read setup. We used cellranger v7.0.1 (10X Genomics) to generate count matrices with default parameters but an optimized transcriptome reference v1.1(*43*) for GRCh38.

### Single nuclei analysis

CellBender v0.3.0(*44*) was used to remove background noise. In CellBender, the number of expected cells and total droplets was based on the UMI curve and the learning rate was reduced based on the automated output report. Further downstream analysis was performed with the R package Seurat v5.0.1(*45*). To increase computational efficiency and reduce memory usage of this large dataset, we used BPCells v0.1.0. Low quality cells were filtered for each sample individually by inspecting quality control plots and removing cells with higher mitochondrial percentages (range: 1-5%), low (<200) or very high molecule counts per cell (range: 6000-9000). Doublets were removed using scDblFinder v1.16.0(*46*) with default parameters. We used Seurat for normalization (LogNormalize, default parameters), identification of highly variable genes (vst method,2000 features), scaling, and performing PCA (default parameters). Next, we used atomic sketch integration (method LeverageScore, 5000 cells) in Seurat to reduce memory usage. Batch effects were accounted for by integrating the samples with scVI v1.0.4(*47*). After inspecting the effect of batch removal, the full dataset was integrated with Seurat. The scVI integrated full dataset was used to calculate the UMAP embeddings with Seurat (30 dimensions). We identified clusters with the FindNeighbors and FindClusters (resolution 0.7) functions. We calculated the top expressing genes of each cluster with FindMarkers (min.pct = 0.1, logfc_threshold = 0.25, p_val_adj < 0.05, two-sided Wilcoxon rank-sum test). To subcluster the immune cells, we selected the immune cell clusters of the main clusters. We then reperformed identification of highly variable genes, scaling, and PCA as explained above. The data were integrated with reciprocal PCA using Seurat and after plausibility checks of the batch removal, UMAP was calculated (30 dimensions). Clusters were identified with FindNeighbors and FindClusters (resolution 2.3). The top expressing genes were identified as explained for the main clusters. Enrichment analysis of top markers (avg_log2FC > 1, p_val_adj < 0.001) was carried out with enrichR 3.2(*48*) with the GO Biological Process 2023 database. Differentially expressed genes between conditions were identified using a pseudobulk method with Libra v1.0.0(*49*) (edgeR 4.0.7(*50*), two-sided likelihood ratio test). Volcano plots of DE genes were created with Enhanced Volcano v1.20.0. Additionally, we determined DE genes in a cluster-independent manner with miloDE v0.0.0.9 (*51*) following the tutorial. Briefly, neighborhoods were assigned based on the scvi integrated data (k = 30, prop = 0.1, d = 30) and DE testing was conducted between conditions (min_count =10). To determine the cluster abundance, we used propeller(*52*) (part of speckle v1.2.0). We used the co-varying neighborhood analysis (CNA), implemented in the rcna package v0.0.99(*53*), to compute correlations between clinical phenotypes or histological measures and our single cell data independent of clustering controlling for age and sex. To classify patients we used the PCA of the cluster abundance and multi-resolution variational interference (MrVI)(*26*). PCA of the cluster abundances and variable contributions were calculated and visualized with FactoMineR v2.9(*54*). To run MrVi v0.2.0, the Seurat object was converted to h5ad with sceasy v0.0.7(*55*) and read as an AnnData object in Python. The MrVI model was trained following the tutorial. The sample-sample distances were predicted with MrVI (get_local_sample_representation) for each cell and then averaged over all cells. Sample-samples distances were visualized in a heatmap using pheatmap v1.0.12 in R (ward.D2,, euclidean distance measure).

### Comparison with published datasets

We downloaded the publicly available annotated data from Yim et al.(*9*) (GEO GSE182098, sciatic nerve), Gerber et al(*10*), (https://snat.ethz.ch/seurat-objects.html, 10X Genomics P60), Mathys et al. (https://compbio.mit.edu/scBBB/ ROSMAP vascular cells)(*14*), and Wolbert et al. (*8*) (GEO GSE142541, mouse). If necessary, the data were preprocessed with Seurat, i.e. normalized with LogNormalize, highly variable genes were identified, scaled, and PCA was performed with default parameters. To identify novel marker genes, cell markers in the published datasets were determined with the FindMarkers function in Seurat (min.pct = 0.1, logfc.threshold = 0.25, p_val_ad < 0.05). To annotate our dataset based on the published dataset, rodent gene names were converted to orthologues using homologene v.1.4.68 (homologeneData2 database). We then classified our cells based on the annotated reference dataset using FindTransferAnchors and TransferData functions in Seurat with default parameters. Cells were labeled as unknown if the Seurat prediction score was below 0.3.

### Histology: Semi-thin sections and analysis

After biopsy, ⅓ of the sural nerve was embedded in Epoxy resin (Fig. 1A, experimental setup). Semithin sections (1 µm) of sural nerves were cut using a EM UC7 ultramicrotome (Leica Microsystems) and stained with toluidine blue. The sections were scanned with a slide scanner (Münster/Würzburg: Grundium Ocus40; Essen: Zeiss Axio Scan Z.1, Hitachi HV F203SCL camera). After blinding, the entire number of total myelinated axons per sural nerve were counted manually by one investigator using the CellCounter plug-in of ImageJ v1.36. Physiologically unmyelinated axons (diameter <1 µm) and Remak-bundle fibers were not included.

The ImageJ g-ratio plugin was used to determine myelin thickness and axonal diameter of 15% of the total myelinated axons per sural nerve. In randomly chosen myelinated axons the inner and outer rim of a myelin sheath were manually traced by one investigator. The g-ratio was calculated by dividing the axonal circumference (i.e., inner rim of the myelin sheath) by the outer circumference of the respective myelin sheath, presuming circular axons. The g-ratio values of each patient were plotted against the calculated axon diameters.

### Preparation of sural nerve cross sections and Xenium spatial transcriptomics

We created a custom stand-alone gene panel for spatial transcriptomics using the commercial Xenium Analyzer platform. The panel was designed based on our single nuclei analysis, mostly focussing on cells that demonstrated a potential role in polyneuropathies, including Schwann cells (mySc, nmSC, repairSC), perineurial cells and vascular cells such as pericytes and endothelial cells. We included a minimum of four top marker genes of each of the clusters and also defined twenty DE genes that were differentially expressed between the different PNP subtypes. We additionally included well known markers of both epi- and endoneurial fibroblasts and different subtypes of leukocytes including B cells, T cells, and macrophages, adding up to a total of 99 genes (Suppl. Tab. 7).

In total, 8 sural nerve formalin-fixed paraffin-embedded (FFPE) samples were included for Xenium spatial transcriptomics: 2 CTRL (S22, S24), 2 CIDP (S01, S11), 2 CIAP (S04, S14) and 2 VN (S29, S30) samples (Suppl. Tab. 1). Samples were primarily selected based on their optimal/intact morphology in the semithin sections and their center with the aim to limit center bias..

Sample preparation and processing was carried out by the CMCB Technology Platform Core Facility EM and Histology and the DRESDEN-concept Genome Center at the Technical University Dresden, in Germany. Samples were processed according to manufacturer protocols (10x protocols, CG000580, CG00582 and CG00584). Briefly, 4 µm cross sections were collected, floated in a 37°C water bath, and adhered to Xenium slides (10x Genomics, PN 1000460). Samples were deparaffinized in 2x Xylene and rehydrated in a descending series from 100% ethanol to MilliQ water. After inserting slides into xenium cassettes, samples were decross-linked and incubated overnight (22 hours) with padlock probes, followed by a post-hybridization wash. Subsequently, padlock probes were ligated, followed by rolling circle amplification, autofluorescence quenching, and nuclear staining (DAPI). Slides were loaded on a 10X Xenium Analyzer (software v. 1.6.1.0), for region selection, with each region corresponding to one entire nerve bundle.

Primary analysis, image processing and decoding, and secondary analysis, cell segmentation and transcript mapping, were performed on-instrument with the analysis software Xenium-1.6.0.8 resulting in a cell-feature matrix and an initial clustering. For cell segmentation, a custom neural network on DAPI images was initially used for nucleus segmentation. Then nucleus boundaries are expanded by 15 µm or until they encounter another cell boundary in X-Y to define cell boundaries. Transcripts were mapped to these 2D shapes according to their X and Y coordinates. Gene expression mapping was visualized using the Xenium Explorer software v.1.3. The post Xenium H&E staining was processed according to the manufacturer protocol, including quencher removal in advance (see 10x Genomics protocol CG000613/A).

#### Xenium data analysis

Xenium data was loaded into Seurat. We removed cells with less than 10 molecules per cell. We performed normalization with SCTransform (default parameters) and computed PCA for each sample. To match our snRNA-seq data to our Xenium data, each cell was classified with the FIndTransferAnchors and TransferData function based on our previously annotated snRNA-seq data (downsampled to 1000 cells per cluster). Cells with a prediction score below 0.3 were labeled as unknown. The 24 main clusters were then aggregated into 8 larger groups (SC, endoC, periC, epiC, VSMC, PC, EC, IC). For manual quantification of endo- vs. epineurial transcripts the Xenium images were loaded into Xenium Explorer 1.3.0 (10x Genomics) and endoneurial areas were manually outlined using the selection tool while using visualization of Schwann cell marker transcripts (*EGR2, NGFR, SOX10, S100B)* to reliably identify such endoneurial areas. The area of each image visibly unoccupied by nerve-associated tissue was selected as epineurium. The area (in μm2), the number of total segmented cells, and the number of immune cell-associated transcripts (leukocytes: *PTPRC*/CD45, B cells: *MS4A1*/CD20, T cells: *CD3E*) was quantified and recorded in each selection area. The density of the respective transcript per total endorial and epineurial area were calculated for each sample. Selected transcripts were visualized in Xenium Explorer 1.3.0 (10x Genomics). Images were integrated with corresponding H&E staining to visualize the spatial orientation.

## Acknowledgements

We are deeply indebted to the patients for their participation. We thank Rebecca Ley from the Institute of Neuropathology Münster for excellent technical histological assistance. We would like to thank Kristina Wagner for the technical assistance with processing of the samples in Essen. This work was supported by the Core EM and Histology, a core facility of the CMCB Technology Platform at the Technische Universität Dresden. We thank Susanne Weiche for excellent technical assistance in preparing the Xenium slides. Part of the calculations were performed on the high performance computing (HPC) cluster PALMA II of the University of Münster, subsidized by the DFG (INST 211/667-1).

## Author contributions

M.H. and G.M.z.H. conceived and supervised the study. M.H., J.T, A.K.M, M.M., M.S., A.-K.U., C.D., K.K., F.S, B.E, K.L., and C.S. were involved in collecting human sural nerves. J.T. performed nuclei extraction and snRNA-seq experiments. I.-N.L. carried out sequencing of snRNA-seq. J.W. planned the Xenium analysis. J.A., A.D. performed Xenium experiments. F.D. determined g-ratios. M.H. and C.S. gathered sample and patient metadata. M.H. performed computational analyses. C.T., R.S., R.F., H.W., C.S., and M.S. co-supervised the study. A.-L.B. designed the figures and schemes. M.H., J.W. and G.M.z.H. wrote the manuscript. All authors read and approved the manuscript.

## Data and code availability

All raw and processed single cell sequencing data with sample and cluster annotations will be publicly available in GEO. An interactive version of the snRNA-seq data, created with cerebroApp v1.4.2(*56*) and an interactive version of the Xenium data, created with TissUUmaps v3.1(*57*), is available at http://pns-atlas.mzhlab.com. The code is publicly available at https://github.com/mihem/pns_atlas.

## Supplementary Materials

Fig S1 to S6 Tables S1 to S9

